# The Overexpression of AmpC is Not Required but May Contribute to Imipenem/Relebactam Non-susceptibility

**DOI:** 10.1101/2023.08.15.553407

**Authors:** Shawn Freed, Nancy D. Hanson

**Affiliations:** Department of Medical Microbiology and Immunology, Creighton University School of Medicine, Omaha, Nebraska, United States of America; Creighton Center for Antimicrobial Resistance and Epidemiology, Creighton University School of Medicine, Omaha, Nebraska, United States of America

## Abstract

In the US, carbapenem resistance in *Pseudomonas aeruginosa* is strongly linked to the regulation of chromosomal resistance determinants, AmpC and OprD. The β-lactamase AmpC requires overexpression and genetic modifications to be capable of inhibiting imipenem activity. The outer membrane porin OprD can be downregulated or undergo genetic modifications that strongly correlate with imipenem non-susceptibility. Co-administration of imipenem and the β-lactamase inhibitor, relebactam, can lower imipenem MICs and restore susceptibility. However, how this occurs in *P. aeruginosa* isolates that do not overproduce AmpC or produce a functional OprD for imipenem entry is not understood. Therefore, we investigated whether imipenem could enter *P. aeruginosa* in the absence of OprD and whether any of the chromosomal β-lactamases (AmpC, OXA-51, PIB-1) contributed to imipenem and/or imipenem/relebactam non-susceptibility. This investigation evaluated 17 imipenem non-susceptible clinical isolates and 3 laboratory strains of PAO1, two of which were porin deletion mutants for either *oprD* or *opdP*. Expression of OXA-50 and PIB-1 RNA was similar to PAO1. However, all 20 isolates exhibited AmpC induction under sub-lethal exposure to imipenem. This occurred in the absence of detectable OprD protein in 18 isolates. Collectively, our data identify that OprD is not the only channel required for imipenem entry and that in many isolates the restored susceptibility to imipenem by imipenem/relebactam was due to the interaction of relebactam on the overexpression of AmpC due to imipenem induction.

## Introduction

*Pseudomonas aeruginosa* harbors chromosomal mechanisms that render antibiotics from multiple classes ineffective^1^. These mechanisms contribute to the high mortality and morbidity that is associated *with P. aerugnosa* infections ^2^. Carbapenem antibiotics are a last-line choice for critical *P. aeruginosa* infections with nearly 13% and 18% of *P. aeruginosa* infections resistant to at least one carbapenem in the US and Europe, respectively^3,4^. Among carbapenem resistant *P. aeruginosa*, only 2% are linked to the presence of a carbapenemase in the US^5^. Therefore, the majority of carbapenem resistant strains of *P. aeruginosa* is largely due to chromosomal mechanisms. Overexpression of the AmpC β-lactamase, efflux pump overexpression, and the downregulation of the OprD outer membrane porin are reported to have the biggest impact on carbapenem non-susceptibility^1,6^.

Imipenem has been shown to evade overexpressed efflux mechanisms and remain a potent option against *P. aeruginosa* efflux mutants^7^. However, *P. aeruginosa* may harbor variant PDC alleles with expanded spectrums of hydrolysis, and coupled with reduced outer membrane permeability lead to emergence of resistance^6^. Sub-lethal concentrations of imipenem can also induce the chromosomal AmpC, which can lead to therapeutic failure for some anti-pseudomonal β-lactams^8^. Relebactam was developed to help restore susceptibility to imipenem when given in combination. Relebactam preferentially binds to a wide range of AmpC (PDC) alleles, preventing them from hydrolyzing or binding to imipenem. Many isolates that are non-susceptible to imipenem alone, have been shown to be susceptible when paired with relebactam^9^.

OprD is an outer membrane channel that selectively allows small peptides and basic amino acids to permeate into the periplasmic space^10^. It has been shown that imipenem translocates through OprD^11^. Modifications in the *oprD* gene can lead to a loss in porin production or confirmational changes to the structure of the porin, both of which can lead to imipenem non-susceptibility^12,13^. OprD has 17 related homologues, 8 of which have substrates experimentally defined^14^. Among these is the porin, OpdP. Genetic analysis revealed OpdP to be the most similar to OprD and *in silico* studies have implicated that imipenem can be translocated through OpdP^15,16^.

We hypothesized that imipenem, when combined with relebactam, was capable of inducing AmpC and that in the absence of OprD, entry of imipenem occurred, resulting in AmpC induction. Therefore, in this study, we investigated the contribution of AmpC on imipenem non-susceptibility and the ability of imipenem to enter the cell in the absence of OprD. The data presented in this study demonstrates that in the absence of OprD, imipenem can enter the cell resulting in the induction of AmpC, thereby identifying that OprD was not the only entry point for imipenem. The ability of AmpC to be induced by imipenem in the absence of OprD provides a substrate that relebactam can act upon thereby reducing the imipenem/relebactam MIC. The induction of AmpC by imipenem regardless of being administered alone or in combination with relebactam indicates that that the overproduction of AmpC contributes to imipenem non-susceptibility.

## Results

Increased susceptibility to imipenem/relebactam compared to imipenem alone, implies that the AmpC β-lactamase can impact imipenem susceptibility (Table 1). However, the contribution of AmpC overproduction to imipenem/relebactam non-susceptibility has not been evaluated. Therefore, we evaluated *ampC* RNA expression in imipenem non-susceptible clinical isolates. The imipenem/relebactam MICs for isolates in this study ranged from 4 to 256 ug/ml (Table 1). Compared to PAO1, *ampC* RNA expression of the clinical isolates ranged from 1 to 296-fold, with 7 isolates (41.2%) having greater than a 5-fold increase (Figure 1A). Four isolates with a greater than 5-fold increase in *ampC* RNA expression remained susceptible to imipenem/relebactam, while 6 isolates had less than a 4-fold increase but imipenem/relebactam MICs were above the resistant breakpoint (Table 1 and Figure 1A). In addition to *ampC*, 2 other chromosomally encoded β-lactamases are present in *P. aeruginosa*: OXA-50 and PIB-1. To determine if these enzymes contributed to the imipenem or imipenem/relebactam MICs, RNA expression of these genes was evaluated. The expression of these chromosomal genes was low. OXA-50 expression ranged from a 3-fold increase to an 11-fold decrease relative to PAO1. PIB-1 ranged from no change to an 11-fold decrease compared to PAO1 (Table 1).

**Figure 1.**
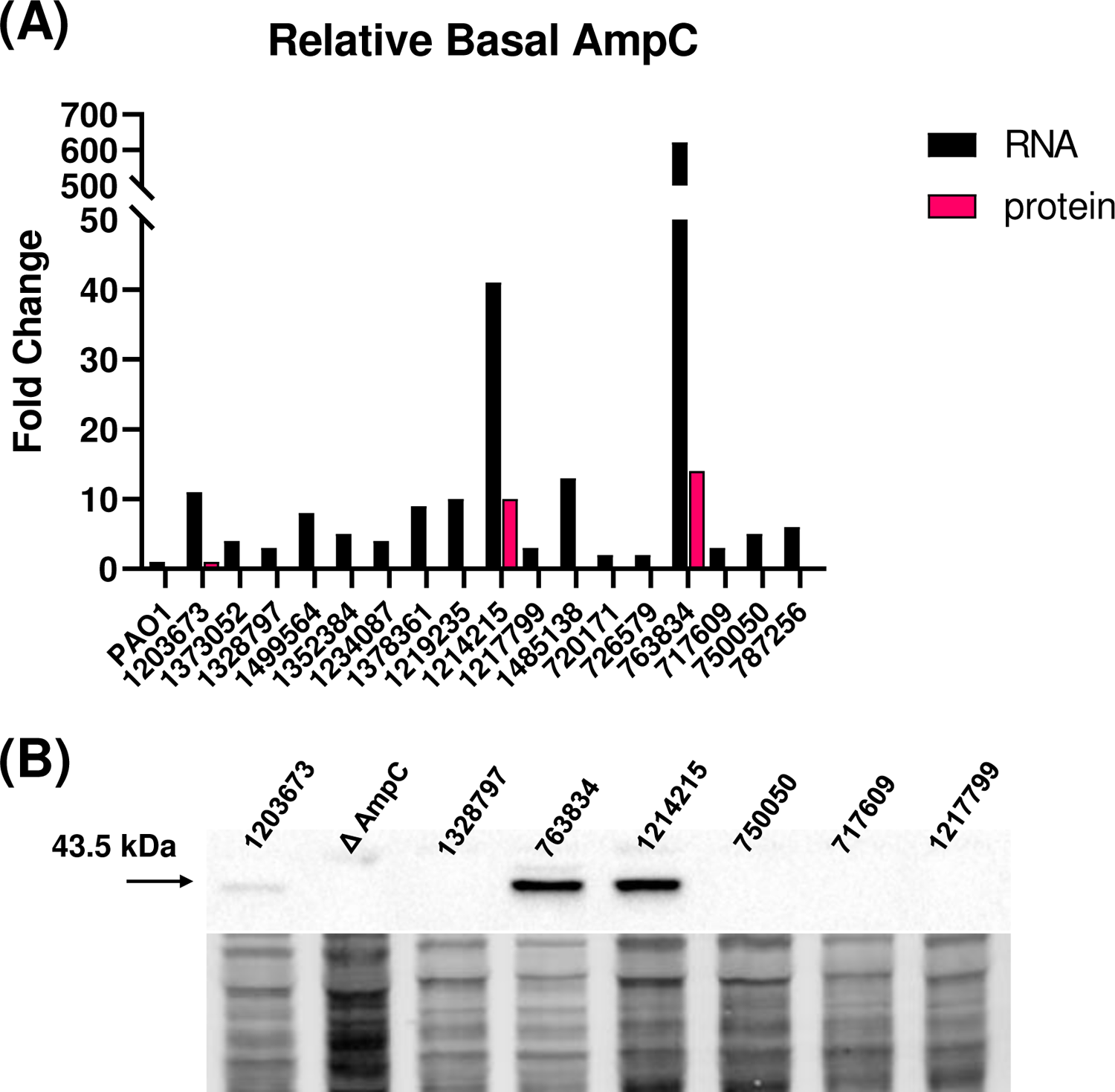
Clinical isolates exhibit increased RNA expression of AmpC, but protein levels do not correlate. (A) Cells were grown to mid-logarithmic phase before being harvested for qRT-PCR (black bars) and western blot (magenta bars) analysis. Fold change in RNA expression was calculated using 2^-ΔΔCT^ method using PAO1 as the comparator isolate. Densitometry analysis from western blot was used to calculate fold change using 1203673 as the comparator isolate. All analysis were conducted in triplicate, using fresh cultures each time. Coefficient of variance was <10%. (B) Representative western blot showing distinct bands for 1203673, 1214215, and 763834. Total protein image is used as a loading control and normalization factor. All experiments were conducted in triplicate, using fresh cultures each time.

**Table 1.** Imipenem MICs and RNA expression of resistance mechanisms for *Pseudomonas aeruginosa* clinical isolates and lab strains. ^a^ Strains ΔOprD and ΔOpdP obtained from Manoil Transposon library^26^. Clinical isolates were obtained from Merck. ^b^ Susceptibilities of clinical isolates were obtained through Merck from IMHA and confirmed locally via E-test. Lab strain susceptibilities obtained via E-test. Phenotypic interpretation: S, susceptible; I, intermediate; R, resistant. Imi, imipenem; Imi/Rel, imipenem/relebactam. ^c^ Relative RNA expression levels of 3 chromosomal β-lactamase enzymes (AmpC, Oxa-50, Pib-1) and 2 porins (OprD, OpdP) determined using qRT-PCR and fold-change calculated using 2^-ΔΔCT^ method with PAO1 as the comparator. Fold change values rounded to the nearest whole-number. ND, not detected after 40 amplification cylces.

| Strain<sup>a</sup> | MIC in µg/mL<br>(Susceptibility<br>Phenotype)<sup>b</sup> |  |  | RNA fold change<br>(Relative to PAO1)<sup>c</sup> |  |  |  |
| --- | --- | --- | --- | --- | --- | --- | --- |
|  | Imi | Imi/Rel | AmpC | OprD | OpdP | Oxa-50 | Pib-1 |
| PAO1 | 0.25 (S) | 0.5 (S) | 1 | 1 | 1 | 1 | 1 |
| 1203673 | 4 (I) | 0.12 (S) | 7 | -1742 | 1 | 1 | -1 |
| 1373052 | 4 (I) | 0.5 (S) | 2 | -340 | 14 | -11 | -6 |
| 1328797 | 4 (I) | 2 (S) | 2 | -2 | 8 | -1 | -2 |
| 1499564 | 8 (R) | 0.25 (S) | 5 | 4 | 6 | 1 | -11 |
| 1352384 | 8 (R) | 1 (S) | 3 | -31350 | 4 | -1 | -7 |
| 1234087 | 8 (R) | 8 (R) | 3 | -4 | 6 | 3 | -5 |
| 1378361 | 16 (R) | 0.25 (S) | 6 | ND | ND | 2 | -4 |
| 1219235 | 16 (R) | 2 (S) | 6 | -64092 | 2 | 2 | -4 |
| 1214215 | 16 (R) | 16 (R) | 26 | 1 | 10 | 1 | -5 |
| 1217799 | 32 (R) | 0.5 (S) | 2 | -28 | 20 | -2 | -7 |
| 1485138 | 32 (R) | 4 (I) | 8 | -18814 | 12 | 2 | -4 |
| 720171 | 32 (R) | 32 (R) | 2 | -106 | 22 | 2 | -9 |
| 726579 | 64 (R) | 8 (R) | 1 | -24146 | 269 | -2 | -5 |
| 763834 | 64 (R) | 16 (R) | 296 | -4 | 4 | -2 | -5 |
| 717609 | 64 (R) | 32 (R) | 2 | -114 | 6 | 1 | -2 |
| 750050 | 128 (R) | >32 (R) | 3 | ND | 123 | -4 | -1 |
| 787256 | 256 (R) | >32 (R) | 4 | ND | 3 | 2 | -3 |
| ΔOprD | 32 (R) | 4 (I) |  |  |  |  |  |
| ΔOpdP | 2 (S) | 0.5 (S) |  |  |  |  |  |

As RNA expression does not always correlate with protein production, Western blots using a polyclonal antibody directed toward the entire amino acid sequence of AmpC was used to determine the relative protein levels of AmpC in the clinical isolates compared to PAO1 (Figure 1B). Only 3 isolates (17.6%): 1203673, 1214215, and 763834, had detectable levels of AmpC protein. Isolates 1214215 and 763834 exhibited the highest RNA expression of 41- and 622-fold respectively compared to PAO1. Both isolates were resistant to imipenem and imipenem/relebactam (Figure 1A). In contrast, 1203673 which was intermediate (MIC = 4) to imipenem alone and susceptible to imipenem/relebactam had only a 7-fold increase in *ampC* expression relative to PAO1 (Figure 1A). Interestingly, 4 isolates (23.5%), 1499564, 1378361, 1219235, and 1485138, had RNA expression between 5 and 8-fold higher than PAO1 but had no detectable AmpC protein (Figure 1A and B). These 4 isolates were resistant to imipenem (MIC = 8 to 32 µg/ml) but 3 of those isolates (1499564, 1214215, and 1219235) were susceptible to imi/rel (MIC = 0.25 to 2 ug/ml) (Table 1). Isolate 1485138 had an intermediate imipenem/relebactam MIC of 4 ug/ml. These data suggested overexpression of *ampC* was not necessary for imipenem/relebactam non-susceptibility, but it may contribute to a non-susceptible phenotype in combination with other mechanisms. However, modifications in genes associated with *ampC* expression or modifications in the *ampC* structural gene could impact the expression and therefore the imipenem/relebactam non-susceptibility. Therefore, we evaluated whole genome sequencing data for mutations that might impact the RNA expression levels observed.

Chromosomal AmpC can influence the imipenem MIC through overproduction of the enzyme in conjunction with hydrolytic properties associated with allelic variants of the enzyme^17–20^. Pseudomonas-derived cephalosporinase (PDC) alleles containing an alanine substitution at residue 105 in addition to the overexpression of the enzyme may contribute to imipenem non-susceptibility^6^. We used whole-genome sequencing to identify the PDC allele harbored by each clinical isolate and any mutations in genes associated with the induction pathway for AmpC, including mutations within AmpD, AmpR, NagZ, PBP-4 and SltB1^21^. Nine different PDC variants were identified in the 17 isolates with PDC-3 present in 7/17 isolates. PDC alleles encoding the T105A substitution were present in 15/17 of those isolates with isolates 1328797 and 1378361 having a wild-type codon at position 105 (Table 2). Although there were a few mutations in genes associated with the induction pathway only two of the isolates (1214215 and 763384) over expressed AmpC both at RNA and protein levels compared to PAO1. These two isolates were resistant to imipenem and resistance was not alleviated by the addition of relebactam. However, all isolates except 1203673, 1373052 and 1234087 had mutations in AmpD and 1373052, 1328797, 1219235, 1485138, and 717609 had mutations within AmpR and yet none of these isolates overexpressed AmpC (Table 2). These data suggested that modifications in the structural gene of AmpC nor any pathways leading to derepression were responsible for imipenem/relebactam non-susceptibility.

**Table 2.**
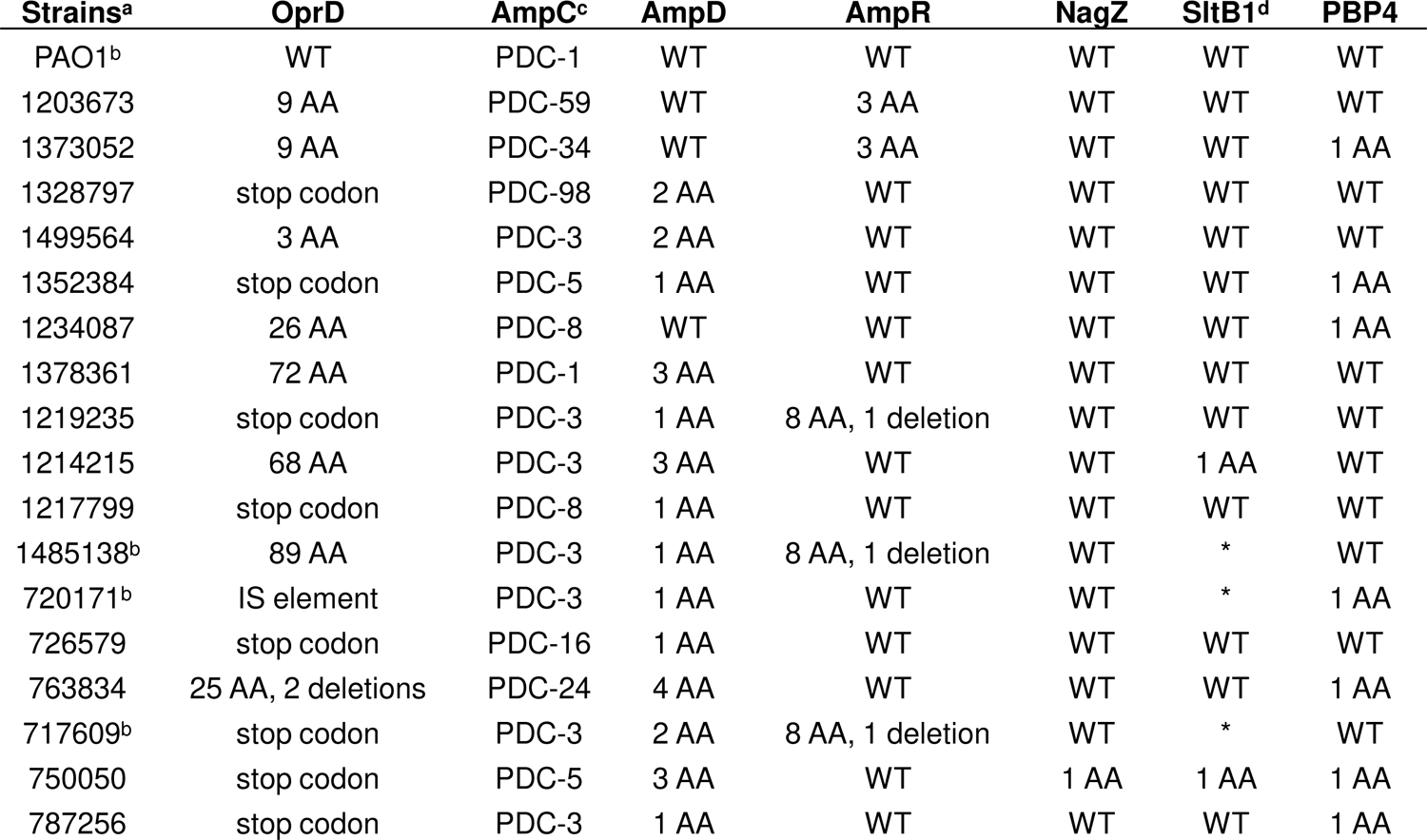
Molecular characterization of resistance mechanisms and induction pathways in clinical isolates of *Pseudomonas aeruginosa*. ^a^ Isolates were sequenced using Illumina MiSeq unless noted otherwise. ^b^ Isolates sequenced by SeqCenter ^c^ AmpC PDC alleles provided by Merck ^d^ 3 sequences could not be retrieved from SeqCenter analysis, denoted by *

Loss of OprD has been shown to play a critical role in emergence of imipenem non-susceptibility^10–12,18,20^. Therefore, we wanted to examine the contribution of OprD to the various susceptibility patterns observed in these isolates by evaluating the RNA and protein production of OprD. All the clinical isolates had reduced *oprD* RNA expression compared to PAO1, with 3 isolates (1378361, 750050, 787256) having no detectable RNA expression (Figure 2A). Five clinical isolates (1328797, 1499564, 1234087, 1214215, 763834) had between no change and a 4-fold decrease relative to PAO1 but the imipenem MICs for these isolates ranged from 4 to 64 ug/ml (Figure 2A and Table 1). Although the RNA levels for *oprD* varied, none of the isolates had detectable levels of OprD using Western blot analysis (Figure 2B). This was expected for the 12 isolates (70.6%) that showed large negative fold-changes in RNA expression, but surprising for the 5 strains (1328797, 1499564, 1234087, 1214215, 763834) with *oprD* RNA expression similar to PAO1 (Figure 2B). The lab strains, PAO1 and ΔOpdP had detectable OprD protein bands and ΔOprD did not (Figure 2B and S.Figure 1A). Our inability to detect OprD protein prompted us to analyze the sequences for mutations, either within the epitope used to generate the polyclonal antibody or mutations that would truncate the protein.

**Figure 2.**
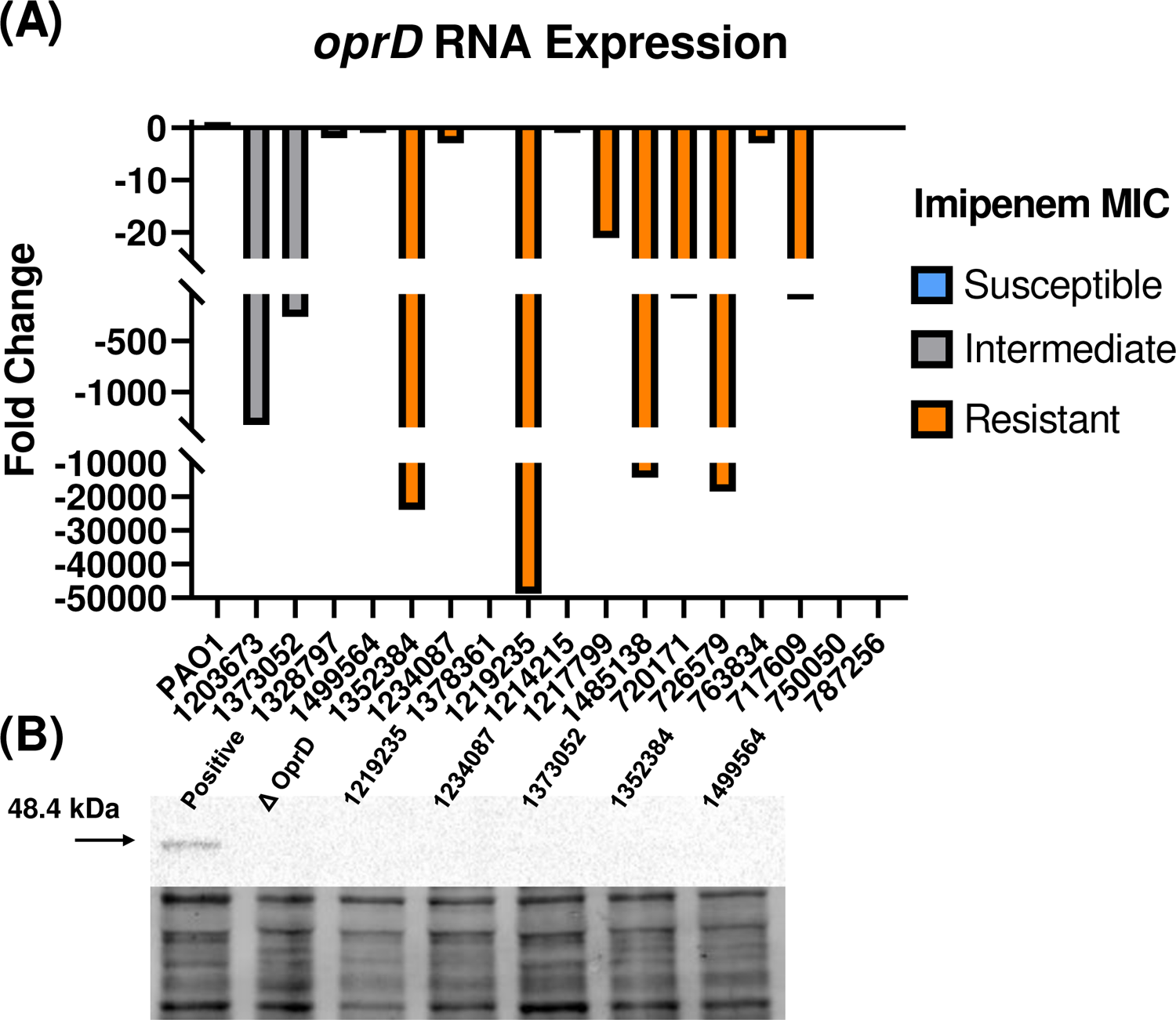
Decreased OpdP expression and production correlates with non-susceptibility to imipenem. (A) Cells were grown to mid-logarithmic phase before being harvested for qRT-PCR and fold change was calculated using 2^-ΔΔCT^ method using PAO1 as the comparator isolate. Imipenem susceptibility phenotypes are indicated by color (grey = intermediate, orange = resistant). 3 isolates (1378361, 750050, and 787256) exhibited no amplification after 40 cycles. (B) Cells grown as stated above. Representative western blot showing lack of protein detection for all clinical isolates. Total protein image is used as a loading control and normalization factor. Densitometry analysis was not performed. All experiments were conducted in triplicate, using fresh cultures each time.

Every isolate harbored amino acid changes compared to PAO1. Nine isolates (52.9%) had a premature stop codon or an IS element that truncated the sequence (Table 2). The remaining isolates were divided into 2 groups: 3 isolates (17.6%) with less than 10 mutations (3 – 9 AA) and 5 isolates (29.4%) with greater than 10 mutations (25 – 89 AA) (Table 2). Isolates with less than 10 amino acid changes had lower imipenem and imipenem/relebactam MICs compared to those with greater than 10 alterations or those with premature stop codons, despite undetectable OprD protein and varying RNA expression (Table 1 and 2). However, despite most isolates (82.4%) having 25 or more amino acid changes or truncations in OprD, relebactam was still able to lower the MIC in 11 (64.7%) of these isolates. These data suggested imipenem penetrated the cell and that relebactam acted upon its target, AmpC.

### AmpC induction by imipenem occurs in absence of OprD

Although the majority of the isolates did not produce detectable levels of AmpC using western blot analysis, treatment of the isolates with imipenem/relebactam lowered the imipenem MICs for 12 of 17 (70.6%) clinical isolates tested (Table 1). Therefore, we investigated whether sub-lethal concentrations of imipenem would induce *ampC* RNA expression and protein production in the clinical isolates.

Induction of AmpC occurred for all the isolates tested (Figure 3A). RNA induction among the isolates varied dramatically, but this was due, in part, to the basal level of *ampC* expression for each isolate. Six isolates (1373052, 1499564, 720171, 726579, 717609, 787256) had greater than 1000-fold increase in RNA expression. This large increase in expression in the presence of imipenem correlated with a low *ampC* basal level of expression with basal level CTs greater than 30, apart from isolates 1373052 and 1499564 which had basal CTs of ∼29 and 28, respectively (Figure 3A). 1203673, 1214215, and 763834, isolates had higher levels of basal *ampC* RNA expression (CTs of 26 and 21) and detectable protein but, still exhibited induction by impenem of 254-, 214-, and 13-fold, respectively (Figure 3A). 1352384 and 750050 showed very little induction of RNA, 34- and 16-fold respectively, despite having low basal levels (CTs 28 to 30) (Figure 3A). These variations observed in the induction potential suggested modifications within the induction pathway. Therefore, sequence data was obtained for all isolates (Table 2). 1352384, had no unique mutations within *ampR, ampD,* nor *dacB* (PBP4) (Table 2). PBP4 did harbor a A394P mutation in 1352384, but this residue was also shared with 4 other isolates showing high levels of *ampC* RNA induction, including the derepressed mutant 763834 (Figure 3A). 750050 harbored a unique A107S mutation within NagZ and a unique D283E mutation within the sequence for the lytic transglycosylase, SltB1 (Table 2). These mutations could have contributed to the lower *ampC* induction potential (16-fold) observed. The lab strains: PAO1, ΔOprD, and ΔOpdP, were also examined. These isolates all exhibited different levels of *ampC* RNA induction (42 to 1600-fold) (Figure 3A). All 17 clinical isolates and 3 lab strains had detectable AmpC using Western analysis following induction (Figure 3B). These data taken together, suggest that imipenem entered the cell in the absence of OprD, which induced AmpC production providing the substrate for relebactam. (Table 1, Figures 2B and 3B).

**Figure 3.**
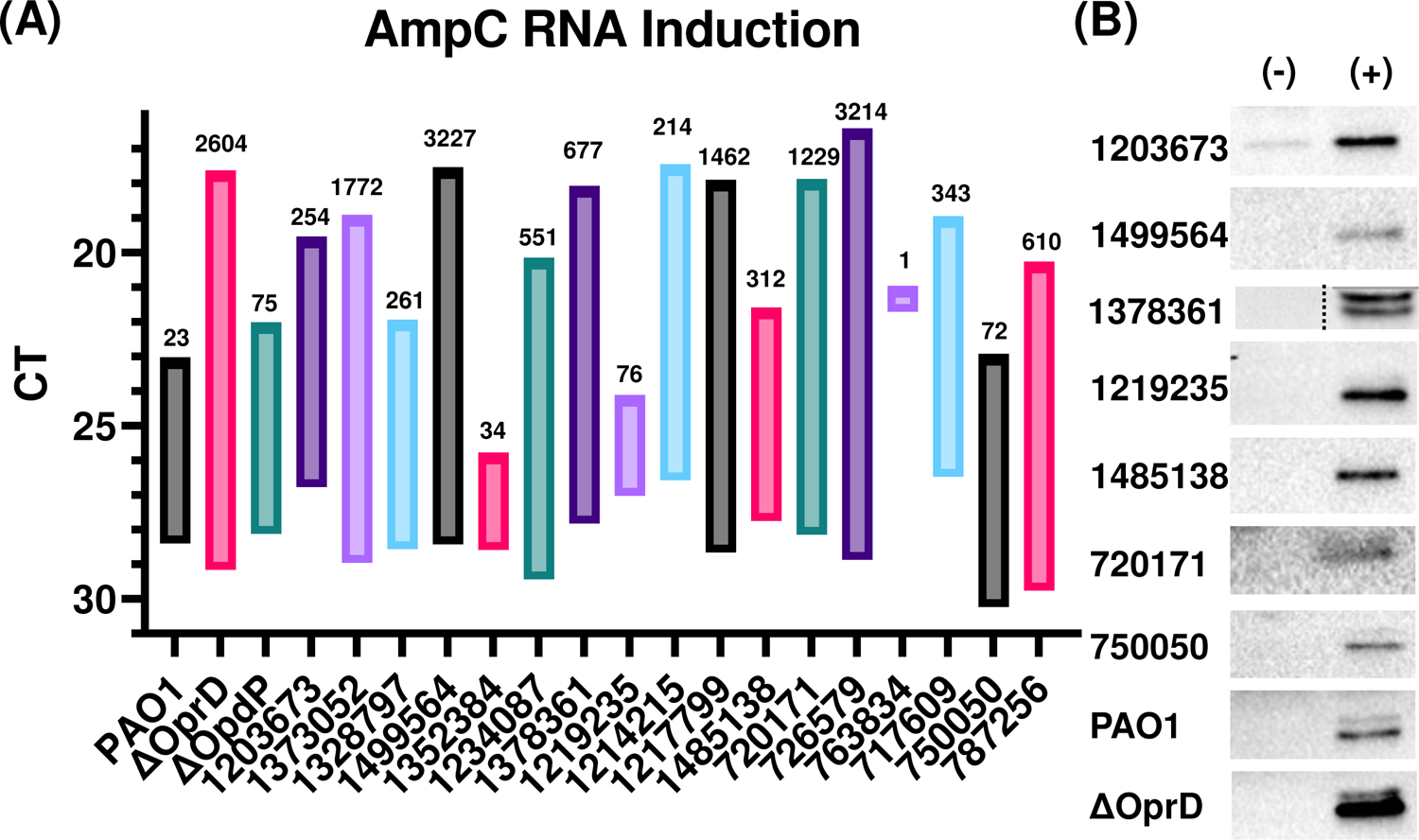
AmpC induction of RNA and protein occurs upon imipenem treatment. (A) Isolates were grown to mid-logarithmic phase then treated with 1/4^th^ their respective imipenem MIC for 15 minutes and subsequently harvested for qRT-PCR and fold change was calculated using 2^-ΔΔCT^ method using untreated respective samples as a comparator. The lower bound of the column represents the basal cycle threshold (CT) for *ampC* and the upper bound is the induced CT. The data label indicates the fold change in expression. (B) Western blot samples were harvested as above. Representative blots show induction of protein following treatment (‘-’ untreated, ‘+’ treated). All experiments were conducted in triplicate, using fresh cultures each time.

### OpdP may contribute to imipenem/relebactam uptake, but is not essential

Previously published studies have suggested an alternate porin within the OprD (OccD) family, OpdP, may be capable of translocating imipenem into the periplasmic space of *P. aeruginosa*^16,22,23^. OpdP expression may compensate for the loss of functionality in OprD seen frequently in *P. aeruginosa* pathogenic strains. We investigated whether the clinical isolates in this study had increased *opdP* RNA expression compared to PAO1. Expression of *opdP* was observed in 16 (94.1%) clinical isolates with 1378361 being the only exception (Figure 4). Of these, 15 (88.2%) isolates had increased relative *opdP* expression compared to PAO1, while 1203673 showed the same level as PAO1 (Figure 4). Despite this, all 16 isolates had CT values for *opdP* (36 to 28), suggesting that OpdP was being produced albeit at low levels in these organisms in the absence of OprD Therefore, OpdP may provide a point of entry for imipenem which could explain the imipenem MICs observed. However, 1378361 did not show any RNA expression for *opdP* nor *oprD*, yet surprisingly AmpC was inducible in the presence of imipenem (Figures 3 and 4). These data imply that another porin(s), in the absence of OprD and/or OpdP, may also be capable of allowing imipenem into the cell.

**Figure 4.**
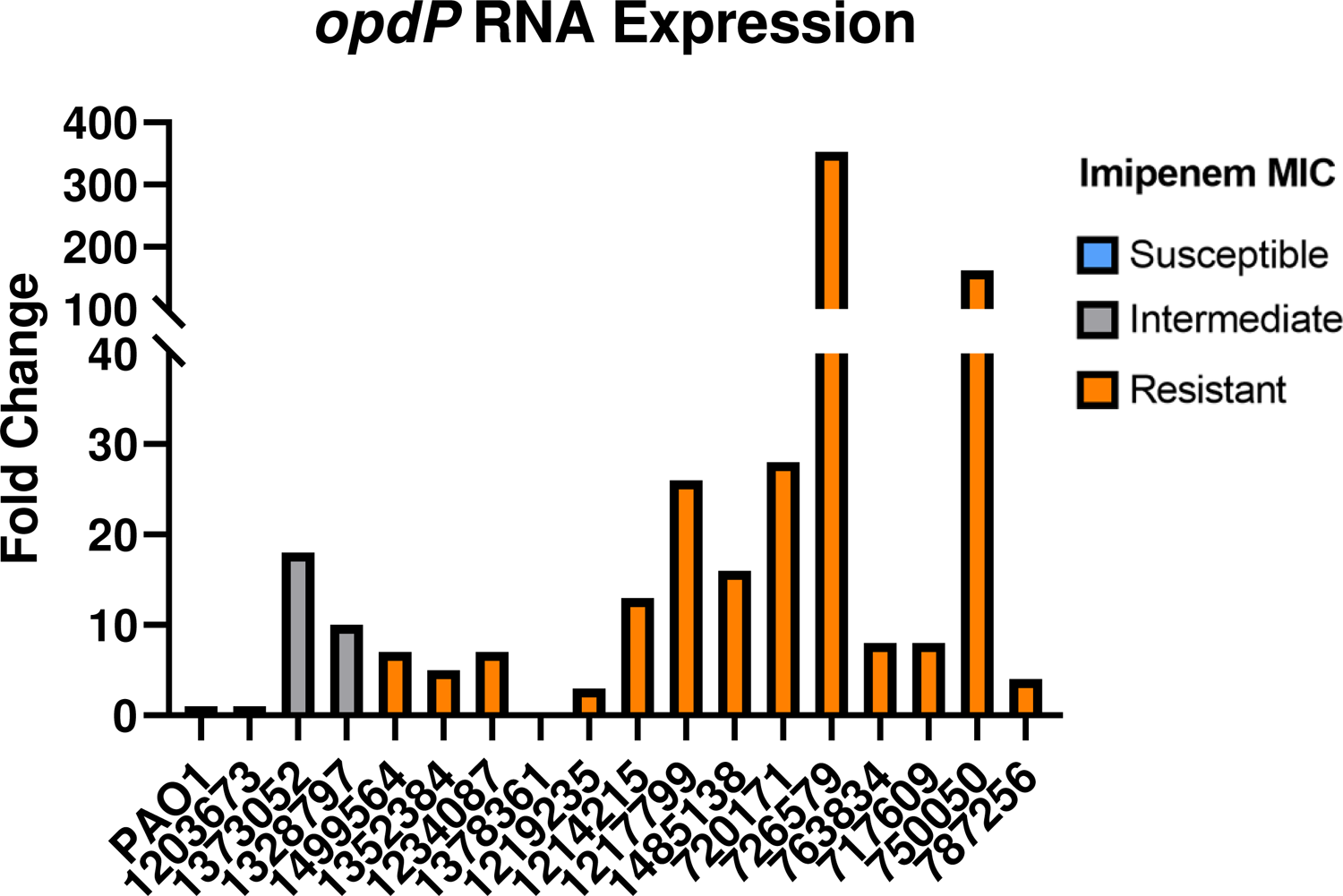
Clinical isolates exhibit *opdP* expression at varying levels. Cells were grown to mid-logarithmic phase before being harvested for qRT-PCR and fold change was calculated using 2^-ΔΔCT^ method using PAO1 as the comparator isolate. Imipenem susceptibility phenotypes are indicated by color (grey = intermediate, orange = resistant). 1378361 exhibited no amplification after 40 cycles. All experiments were conducted in triplicate, using fresh cultures each time.

### RNAseq analysis of isolates under imipenem challenge

We investigated the transcriptome of the 3 lab strains: PAO1, ΔOprD, and ΔOprP, as well as 3 clinical isolates selected for their unique phenotypes. Isolate 1203673 had detectable AmpC protein, downregulated *oprD* expression (1742-fold decrease), but a low imipenem MIC (4μg/mL). Isolate 1378361 had no RNA expression of *oprD* and *opdP*, yet AmpC was still inducible by imipenem. Isolate 1499564 was resistant to imipenem alone (8μg/mL) yet susceptible to all other antibiotics tested. This isolate also had similar expression of both *ampC* and *oprD* compared to PAO1. Principal-component analysis (PCA) was performed for isolates in the absence of imipenem (Figure 5A). Evaluation of 5853 genes showed that transposon mutants deficient in one of the two porins clustered more closely with clinical isolates than with WT PAO1 (Figure 5A). The clinical isolate 1203673 clustered closely with PAO1, which was not surprising given several genotypic similarities in RNA expression as well as lower imipenem MICs (Table 1). Data from the 1499564 isolate replicates clustered the furthest from themselves while ΔOprD clustered the furthest from any of the isolates. These data were interesting and further analysis of the transcriptome related to PAO1 was carried out (Figure 5B).

**Figure 5.**
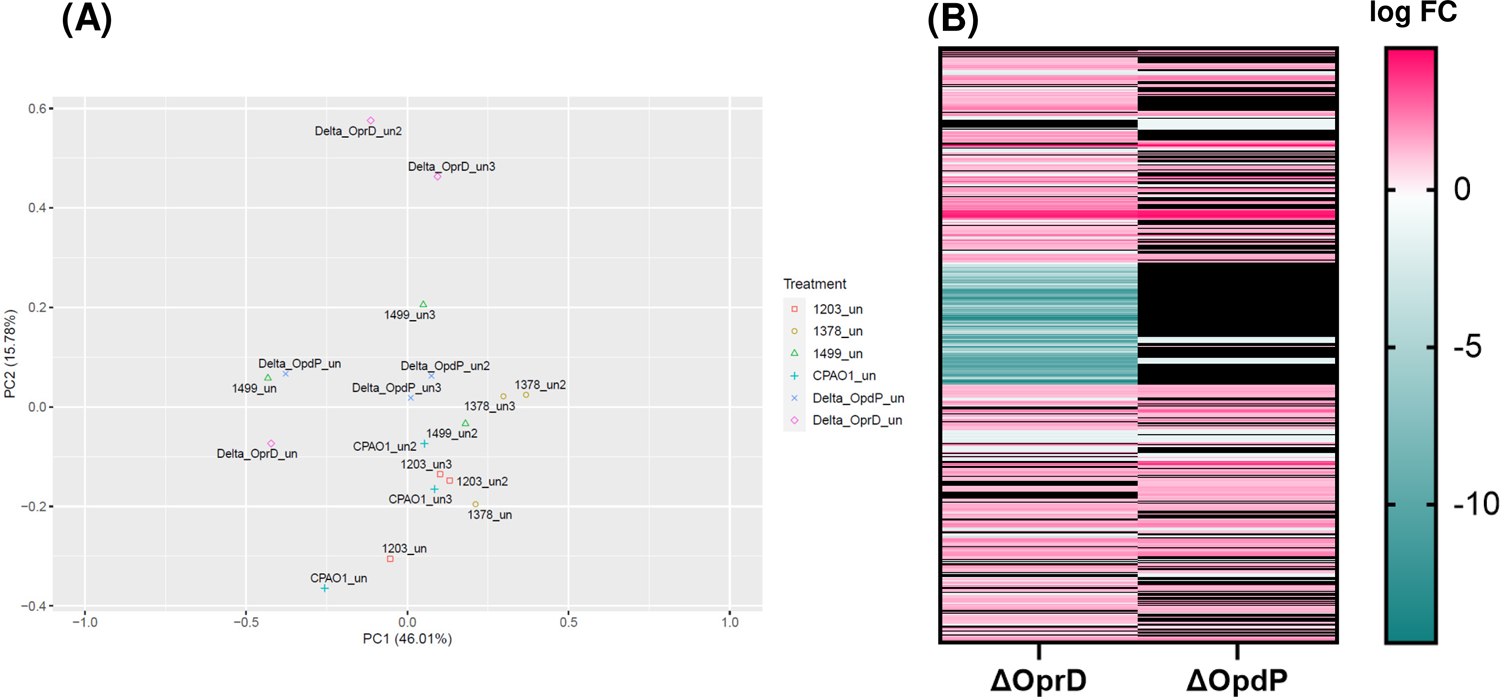
Transcriptomic analysis of 3 *Pseudomonas aeruginosa* laboratory strains and 3 clinical isolates in the absence of imipenem. (A) Principal-component Analysis considering all quantified genes. Each isolate is represented by a different symbol and color (see key). Biological replicates are labeled* within the graph with the replicate number and designated as ‘uninduced.’ (B) Heatmap showing differentially expressed genes of the ΔOprD and ΔOpdP transposon mutants compared to PAO1. log fold-change (FC) is indicated by the colors shown in the key. Blacked out genes are not significantly differentially expressed from PAO1 (*P* < 0.05, log FC > │1│). All experiments were conducted in triplicate, using fresh cultures each time. *A labelling error for ΔOprD_un2 was corrected during analysis.

Comparative analysis between ΔOprD and PAO1 revealed a large contiguous set of 145 genes (*czcC*/PA2522 – PA2377) deleted from the ΔOprD genome. (Figure 5B, Supplemental File 3). Many of these genes were hypothetical, but interestingly porins *opdT* and *opdJ* were deleted. Coupled with *oprD*, this isolate was deficient in 3 OprD-related porins.

Differential RNA expression was determined for 6 isolates (PAO1, ΔOprD, ΔOprP, 1203673, 1378361, and 1499564) with and without imipenem challenge (Figures 6A and 6B). Significance for gene enrichment or depletion was based on the thresholds *P* < 0.05 and log FC > │1│. PAO1 and 1378361 underwent the fewest changes, with only a total of 15 differentially expressed genes (DEGs), with 0 and 3 uniquely differentially expressed in the organism, respectively. Isolate 1378361 did not express either *oprD* or *opdP* by qRTPCR (Table 1). The clinical isolate 1203673 had only 17 significant DEGs and only 5 of them were unique. Clinical isolate, 1499564, was evaluated by RNAseq because it had increased expression of both *oprD* and *opdP* compared to PAO1. RNAseq analysis revealed 142 DEGs and 103 of them were unique. Both transposon mutants were analyzed: ΔOpdP had 52 DEGs and 18 of them were unique, whereas ΔOprD had 139 total and 85 unique. Among all 6 isolates only 3 DEGs were shared: *creD*, *ampC*, and PA4112, and all 3 were enriched (Figure 6A and 6B).

**Figure 6.**
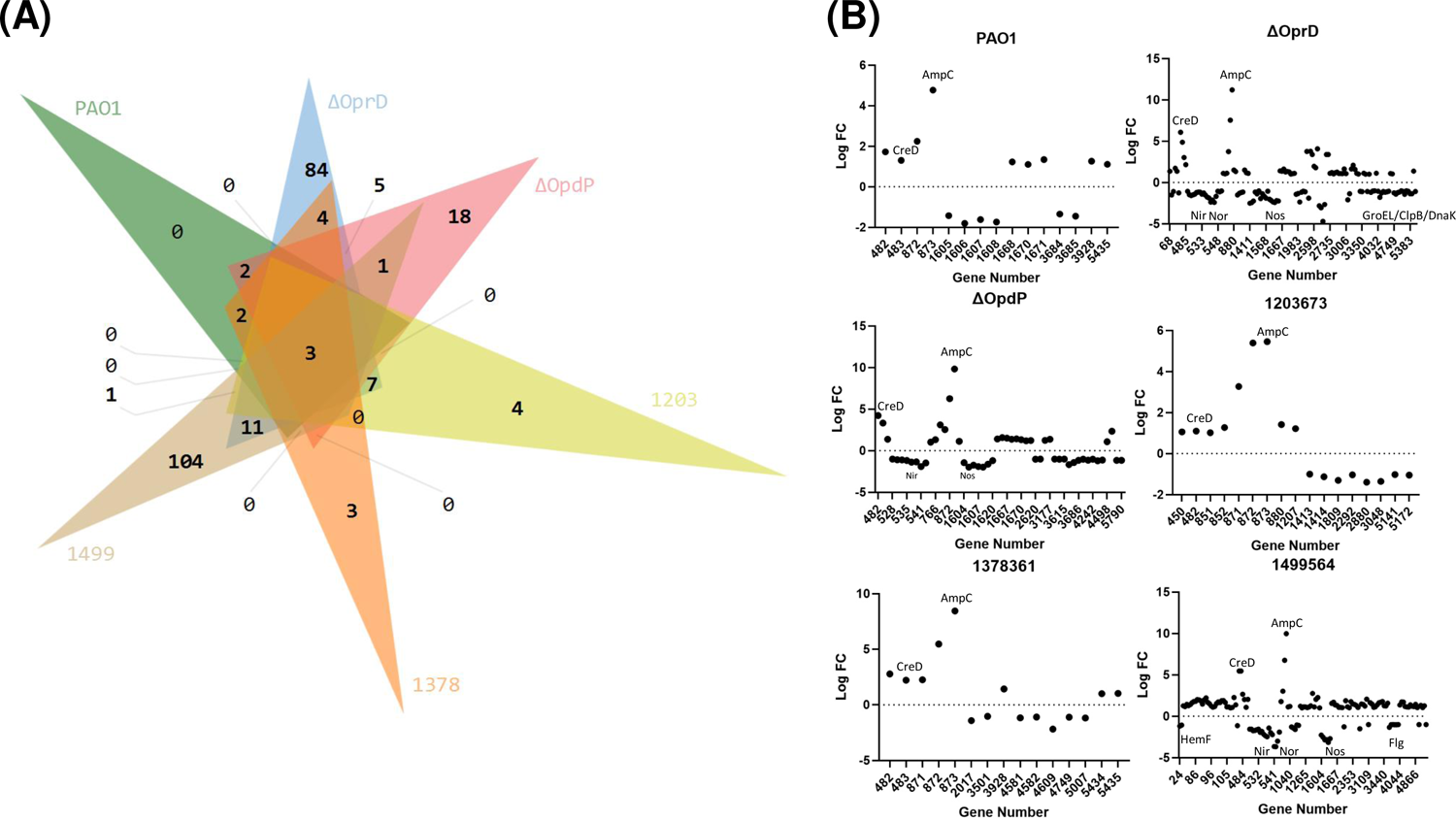
Transcriptomic analysis of *Pseudomonas aeruginosa* isolates treated with sub-lethal concentrations of imipenem. (A) Venn diagram showing significantly (*P* < 0.05) differentially expressed genes that meet a threshold of log fold-change (FC) > │1│Generated using InteractiVenn ^32^. Comparative analysis for lab strains: PAO1, ΔOprD, and ΔOpdP, and clinical isolates: 1203673, 1378361, and 1499564. (B) Differentially expressed genes under imipenem challenge for the isolates PAO1, ΔOprD, ΔOpdP,1203673, 1378361, and 1499564. 1203 = 1203673, 1378 = 1378361, 1499 = 1499564. *Pseudomonas* Genome Database gene annotation numbers are used^33^. CreD, AmpC, GroEL/DnaK/ClpB indicate the position of the respective gene within the differentially expressed sample set. Nir, Nor, Nos, Flg, HemF indicate the position of the respective operons within the differentially expressed sample set. All experiments were conducted in triplicate, using fresh cultures each time.

Between all the isolates, members of the operons for nitric oxide, nitrite, and nitrate reduction were commonly depleted. Nitric oxide reduction and nitrite reductase operons were depleted between log FC −2 and log FC −4 in 1499564 and ΔOprD. (Figure 6B). A nitrous oxide operon was found to be depleted in 1499564, ΔOprD, and ΔOpdP between log FC −2 and −3 (Figure 6B). Isolate 1499564 also had depletions within a flagellar operon and a putative coproporphyrinogen oxidase operon (Figure 6B). Isolate 1378361 had Log FC 1 depletions in genes encoding the chaperon proteins DnaK and GroEL. These depletions were shared only with the ΔOprD isolate (Figure 6B). Many of the DEGs within all 5 isolates were not annotated or were indicated as hypothetical proteins, which poses a challenge for analysis on the changes under imipenem challenge. All DEGs observed for these isolates can be found in supplemental data table 4.

## Discussion

Relebactam was designed to be paired with imipenem as an inhibitor for β-lactamases including the chromosomal AmpC of *P. aeruginosa*. In many cases, *P. aeruginosa* isolates resistant to imipenem alone have decreased and often susceptible imipenem/relebactam MICs. However, the major mechanism of resistance to imipenem is the downregulation of the porin, OprD^17,18,20^. How then, does relebactam help to lower the imipenem MICs when decreased production of the porin OprD is the mechanism attributed to imipenem resistance? It has typically been recognized that overproduction of AmpC does not play a role in imipenem resistance unless modifications in the hydrolytic properties of AmpC accompany the overexpression of the enzyme^12^. Modifications associated with AmpC alleles (PDCs), especially T105A, are capable of imipenem hydrolysis^6^ when AmpC is overexpressed. However, even in the absence of this amino acid modification, isolates were non-susceptible to imipenem. (Table 1). These data strongly suggested that the AmpC alleles present these isolates did not contribute to the imipenem non-susceptible MICs and is supported by the findings of Berrazeg, et al^24^. Imipenem non-susceptibility in the isolates evaluated in this study was attributable to the lack of OprD. But these data did not provide an explanation as to how relebactam lowered imipenem MICs in 71% of clinical isolates tested.

Imipenem induces AmpC expression in *P. aeruginosa*^8^. In the presence of sublethal concentrations of imipenem all the clinical isolates were induced and expressed AmpC (Figure 3). All but 5 of these isolates showed a decrease in imipenem MICs in the presence of relebactam (Table 1). These data suggested that AmpC plays a role in imipenem susceptibility when the enzyme is overproduced. However, the induction potential of the individual clinical isolates was not equivalent. Two isolates, 1214215 and 763834, were derepressed mutants as the basel level of RNA and protein expression was much higher than the other clinical isolates (Figure 1). The imipenem MIC decreased in 763834 when relebactam was introduced but remained above the resistant breakpoint for both organisms (Table 1). Given the difference in the induction potential of these isolates, WGS was used to identify the presence or absence of mutations within genes in the AmpC induction pathway. Mutations or disruptions in this pathway were not identified by whole genome sequencing and could not account for the variation we observed in AmpC induction (Table 2). Variations observed in the levels of induction are directly impacted by the basal level of AmpC expression. All three isolates with detectable AmpC protein (1203673, 1214215, 763834) exhibited comparatively smaller (<260 fold) increases in *ampC* RNA when treated with imipenem (Figure 3A). We tested 2 other chromosomal β-lactamases, OXA-50 and PIB-1, and observed little difference in the RNA expression from PAO1, suggesting they played no role in the imipenem MICs (Table 1).

It is possible that subtle differences in OprD production may limit the amount of imipenem entering the cell and attenuate the induction response in some isolates. It was clear from the WGS data that some of these isolates did not have a functional OprD due to truncations or insertions withing the gene. Most of the OprD protein sequences had premature stop codons and were truncated (Table 2). The remaining isolates harbored multiple mutations that may lead to misfolding and premature degradation (25 or more residue changes) or may be responsible for reduced uptake efficiency (10 or less residue changes). How then, did induction by imipenem occur in absence of a functional OprD. The data presented in this study suggests that imipenem is transported into the periplasmic space through an alternative porin or porins. A PAO1 knock-out strain missing the genes for *oprD* confirmed imipenem entrance by exhibiting AmpC induction in the presence of sublethal concentrations of imipenem (Figure 3B)

Recent literature has indicated that the outer membrane porin OpdP may also allow entry of imipenem.^15,16^. Therefore, we evaluated RNA expression of *opdP*. Our data indicated stable yet low expression of OpdP RNA in almost all clinical isolates (Figure 4). While whole-genome sequence analysis revealed a myriad of mutations within *oprD* sequences compared to PAO1, very few alterations were observed within *opdP.* Whole genome sequence analysis of OpdP showed very few mutations across all isolates, thus indicating a functional porin. Interestingly, an isolate (1378361) was observed that did not have detectable RNA amplification of *oprD* or *opdP*. The isolate exhibited OprD β-barrel structural changes as well as receptor alterations using structural modeling (data not shown), but the OpdP amino acid sequence was identical to PAO1. Despite lacking detectable porins OprD and OpdP, AmpC induction occurred in the presence of imipenem.

Transcriptomic analysis of isolates under sublethal imipenem challenge indicated a decrease in several operons related to nitrite metabolism, nitric and nitrous oxide metabolism, flagellar proteins, and several heat shock protein-related chaperon proteins (Figure 6B). This suggests *P. aeruginosa* undergoes a shift away from metabolic and motile processes when encountering a compound that compromises peptidoglycan integrity (imipenem). CreD and expectedly, AmpC were both enriched under imipenem challenge. CreD has been reported to act as a global virulence regulator and play a role in AmpC regulation ^25^. Unfortunately, many genes that were differentially regulated were not annotated or denoted as hypothetical proteins. Transcriptomic analysis also highlighted the extreme variability between not only clinical isolates, but the laboratory strains. The knock-out strains, generated from PAO1 lineage, differed greatly from the PAO1 used in analysis. ΔOprD RNAseq analysis revealed 145 sequential genes completely absent from the genome when compared to PAO1, ΔopdP, or the NCBI reference sequence and this was validated with WGS (Figure 5 and Supplemental File 3). Among these genes, was *opdT* and *opdJ*. Taken together with our imipenem induction data, this supports the assertion that additional porins contribute to imipenem translocation. Although entry through these other porins may not be as efficient. This information could be used to modify imipenem structure to increase its affinity for other porins so that entry was not so heavily dependent on OprD.

The data presented in this study indicated that imipenem could enter the cell in the absence of OprD and have provided a rationale behind reduction in imipenem MICs when used in combination with relebactam, through the induction of AmpC. Clearly, not all clinical isolates fit the scenario of relebactam decreasing the imipenem MIC, indicating other mechanisms or combination of mechanisms could play a role in the ability of *P. aeruginosa* to become imipenem resistant. Isolate 1234087 had levels of *oprD* (4-fold decrease) and *ampC* (3-fold increase) RNA close to that of PAO1 however, it was resistant to imipenem and the MIC did not decrease with relebactam (Table 1). Conversely, 1352384 had among the lowest *oprD* RNA levels detected (30,000-fold decrease) and weakest induction of *ampC* RNA following imipenem treatment (34-fold increase). Despite this, the imipenem MIC dropped below the susceptible breakpoint (from 8μg to 1μg) when treated with imipenem/relebactam (Table 1). Currently it is not well understood how relebactam enters the periplasmic space. This reinforces the need to explore other outer membrane proteins to understand their contributions to imipenem non-susceptibility.

## Methods

### Bacterial Strains and Growth conditions

A descriptive list of the strains used in this project are given in Table 1. *P. aeruginosa* PAO1 derived knock-out mutants were obtained from the Manoil Lab transposon mutant library ^26^ and were confirmed via PCR for the presence of the transposon and the absence of the associated gene. Cells were plated from frozen % glycerol stocks upon TSA with 5% Sheep Blood and grown overnight at 37°C. Cells were taken and inoculated in 95mL of cation-adjusted Mueller-Hinton broth at an optical density of .1 (Spectra _OD_600). Isolates were grown in a shaking incubator at 37°C to mid-logarithmic phase, denoted by OD of .5 or 1×10^10^ cells.

### Antibiotic Susceptibility Testing

The MICs forPAO1, ΔOprD, and ΔOprP were determined using Etest and interpreted by CLSI guidelines. The MICs for the clinical isolates were performed by IHMA.

### RT-qPCR

Isolates were grown according to method listed above and harvested at mid-logarithmic phase. RNA extraction was carried out using TRIzol™ Reagent (Invitrogen, ThermoScientific) Precipitation of RNA was performed using isopropanol extraction and centrifugation. TURBO DNase (Invitrogen, ThermoScientific) was used to remove DNA contaminants and the RNA was quantified on an Eon spectrophotometer (BioTek). For the imipenem induction experiments, after growth to an OD_600_ 0.5, cultures were treated with 1/4th the respective MIC concentration of imipenem and grown for 15 minutes prior to total RNA isolation. RT-qPCR was performed using QuantiNova SYBR Green 1 Step kit (Qiagen) with 40 cycles of amplification at 57ᵒC performed on an ABI 7500 (Applied Biosystems). Fold change in RNA levels was determined using the 2^-ΔΔCt^ method^27^ using *sodB* as an endogenous control. All experiments performed in triplicate with independent growth and collection. All primers used for RT-qPCR experiments located in Supplemental file 2.

### Immunoblot

Custom polyclonal antibodies specific for AmpC and OprD were generated by GenScript. The epitopes for each are listed in Supplemental file 2. The linear range of detection for the anti-AmpC and anti-Oprd antibodies as well as Stain-Free^TM^ fluorescence was determined via a dilution series western blot using total protein concentrations ranging from 2.5 to 45 μg (Figure S2 and S3). The AmpC and OprD antibodies were diluted 1:20000 and 1:30000 respectively for immunoblot for all analyses.

Isolates were grown according to method listed above and harvested at mid-logarithmic phase. Whole cell lysates were collected by bead-beating (Next Advance Bullet Blender) and protein concentrations determined using a Bicinchoninic Acid Assay (Pierce BCA Protein Assay, ThermoScientific) on an Eon spectrophotometer (BioTek). Total protein lysates were separated via SDS-PAGE and normalized across the isolates using Stain-Free^TM^ (BioRad) fluorescent signal intensity. Following overnight primary antibody incubation, the secondary antibody was used with a dilution factor of 1:40000. To ensure specificity to OprD protein, ΔOprD was used as a negative control (Figure S3). OprD levels were determined and compared PAO1 via densitometry using Image Lab (BioRad) and Microsoft Excel. All protein experiments were performed in triplicate with independent growth and collection.

### Whole-genome and Transcriptomic sequencing

Following growth, DNA from isolates was extracted using a MagAttract Microbial DNA Kit (Qiagen) and whole-genome sequencing performed either on an Illumina MiSeq or completed by SeqCenter. Following RNA isolation, samples were sent to SeqCenter for transcriptomic sequencing and analysis. Reads were quality checked and adapters trimmed using bcl-convert^28^ and mapped using HISAT2^29^. Reads were quantified using Subread’s featureCounts^30^ and the counts normalized through edgeR’s^31^ Trimmed Mean of M values algorithm. Normalized counts were converted to counts per million and differential expression analysis was undergone using edgeR’s Quasi-Linear F-Test functionality against treatment groups. Genes were determined to be significantly differentially expressed with a │logFC│> 1 and *P* value < 0.05. All RNAseq analyses were performed in triplicate with independent growth and collection.

## Data Availability

Whole-genome assemblies will be uploaded to GenBank NIH sequence database under accession nos. (in progress). Raw Illumina RNA read data will be uploaded to the Gene Expression Omnibus (GEO) database under accession no. (in progress).

## Acknowledgements

Funding provided by Merck, Inc via an investigator-initiated grant.

We thank SeqCenter for generating whole genome sequencing data and transcriptomic data for this project.

---

**Supplemental Figure 1.** OprD antibody affinity and KO confirmation. (A) Cells were grown to mid-logarithmic phase before being harvested for western analysis. Whole cell lysates were loaded, and presence of the protein was determined. PAO1 and ΔOpdP have intact *oprD.* The clinical isolate 750050 has truncated *oprD.* OprD molecular weight is 48.4 kDa (B) Total protein image is used as a loading control and normalization factor. 1: PAO1, 2: 750050, 3: ΔOprD, 4: ΔOpdP.

**Supplemental Figure 2.** AmpC protein linearity curve. (A) Cells were grown to mid-logarithmic phase before being harvested for western analysis. Whole cell lysates were loaded in increasing concentrations then incubated with anti-AmpC rabbit primary antibody at 1:20000 concentration and anti-rabbit goat secondary antibody at 1:40000 concentration. Densitometry analysis from was used to determine linear range of detection. AmpC molecular weight is 43.5 kDa (B) Total protein image was used as a loading control and normalization factor. (C) Signal intensity was plotted against protein concentration and linear regression analysis was used to assess linearity. Protein concentrations: 1: 2.5μg, 2: 5μg, 3: 10μg, 4: 15μg, 5: 20μg, 6: 25μg, 7: 30μg, 8: 40μg, 9: 45μg.

**Supplemental Figure 3.** OprD protein linearity curve. (A) Cells were grown to mid-logarithmic phase before being harvested for western analysis. Whole cell lysates were loaded in increasing concentrations then incubated with anti-OprD rabbit primary antibody at 1:30000 concentration and anti-rabbit goat secondary antibody at 1:40000 concentration. Densitometry analysis from was used to determine linear range of detection. OprD molecular weight is 48.4 kDa (B) Total protein image was used as a loading control and normalization factor. (C) Signal intensity was plotted against protein concentration and linear regression analysis was used to assess linearity. Protein concentrations: 1: 2.5μg, 2: 5μg, 3: 10μg, 4: 15μg, 5: 20μg, 6: 25μg, 7: 30μg, 8: 40μg, 9: 45μg.

